# Macrophage-Derived PDGF-BB and GDF-15 Promote Drug Resistance in KRAS-Mutant Colorectal Cancer

**DOI:** 10.64898/2026.04.27.721111

**Authors:** Björk S. Aston, Hammed A. Badmos, Ross Cagan

## Abstract

Macrophages are abundant in the colorectal tumour microenvironment and can alter drug response. Using mouse *Apc*/*Kras*/*Trp53* (*AKP*) colorectal cancer organoids, we found that macrophages and/or macrophage-conditioned medium reduced sensitivity to the MEK inhibitor trametinib and the pan-RAS inhibitor RMC-6236. In contrast, macrophage-conditioned medium had little effect on regorafenib and increased sensitivity to dabrafenib, suggesting that resistance depends on the inhibitory profile of each drug. Secretome profiling identified PDGF-BB and GDF-15 as candidate mediators. Adding both ligands to organoid medium reproduced much of the conditioned-medium effect, whereas either ligand alone was insufficient. Inhibition of PDGFR or RET partially reduced drug resistance, suggesting that PDGF-BB and GDF-15 likely act through canonical signalling by that additional macrophage-derived signals also contribute. Kinome profiling pointed to increased tyrosine kinase signalling during trametinib treatment, with SRC family kinases emerging as a key downstream node. Consistent with this, SRC inhibition reduced the difference between control and conditioned-medium responses. The multi-kinase inhibitor masitinib—which targets several kinases along this resistance network—strongly restored sensitivity to trametinib and RMC-6236. Together, these data define a macrophage-driven resistance network in KRAS-mutant colorectal cancer organoids and support combined inhibition of RAS-pathway and tyrosine kinase signalling.

## Introduction

Colorectal cancer (CRC) is the second leading cause of cancer deaths globally [1]. Approximately 45% of patients have an activating KRAS mutation (K-CRC), commonly in codon 12 or 13, that results in oncogenesis [2,3]. Targeted therapy options remain limited for most patients with K-CRC. Regorafenib is a multi-kinase inhibitor approved for previously treated metastatic CRC. Direct KRAS-targeted combinations are now available for *KRAS^G12C^* tumours [4,5]; this represents only a minority of K-CRC patients, and has demonstrated limited durability and significant side effects [6,7].

Attempts to drug KRAS and its downstream signalling pathway in CRC have to date yielded limited clinical success, including with drugs that have shown success in other KRAS-driven tumour types [8–10]. For example, the potent MEK inhibitor trametinib failed to provide clinical benefit as a single agent despite promising pre-clinical data [11]. This raises two questions that the present work addresses: what factors promote targeted therapy resistance in KRAS-mutant CRC, and can we circumvent this resistance through a rational multi-targeting approach [12–17]? One important consideration in drug resistance is the immune tumour microenvironment (ITME): targeting the ITME provides an opportunity to manipulate local signals that enhance tumour progression and improve treatment efficacy [18–25].

Macrophages are an especially abundant cell type in the colorectal ITME [26–29]. These myeloid cells perform a broad array of important functions: spanning innate and adaptive immune responses, antigen presentation, phagocytosis, and homeostatic functions such as tissue maintenance and repair [30,31]. In the context of cancer, tumour associated macrophages (TAMs) move dynamically between inflammatory (M1-like) and immunosuppressive (M2-like) roles, and this plasticity can impact drug response [32–39]. Part of this resistance may reflect the plasticity of the macrophage secretome; TAMs prime and promote cancer associated pathways, driving proliferation, extravasation, angiogenesis, evasion from immunosurveillance, alongside other metabolic changes [40–45].

In this work, we provide evidence that macrophages secrete factors, two of which are GDF-15 and PDGF-BB, that together impact drug efficacy in pre-clinical models of KRAS-mutant CRC. GDF-15 is clinically relevant in CRC: high serum levels have been shown to correlate with poor patient prognosis and cachexia [46–53]. In the brain, GDF-15 signals through a receptor complex involving RET and GFRAL [54]. However, the role and identity of GDF-15 receptors relevant to CRC are unknown. PDGF-BB drives tumour proliferation, migration and survival by binding PDGFRαα, PDGFRββ and PDGFRαβ, activating JAK/STAT, AKT, PI3K and RAS pathways [55–58]. Intracellular signalling cascades triggered by these transmembrane proteins are linked through the SRC family kinases (SFKs), which in turn can promote both tumour progression and drug resistance through multiple mechanisms [59–62].

Here we identify a macrophage-driven mechanism of drug resistance, mediated by the cytokines GDF-15 and PDGF-BB. Together these factors drive resistance to trametinib through a coordinated kinase network. We use this information to identify a rational drug combination strategy: inhibiting multiple activated kinases, that contribute to this emergent resistance network, reduces macrophage-mediated drug resistance and slows tumour progression.

## Results

### Macrophages impact drug response in AKP organoids

To explore the impact of immune cells on tumour drug response, we co-cultured (i) mouse *Villin–CreERT2: Apc^fl/fl^ Kras^G12D/+^ Trp53^fl/fl^* (*AKP*) organoids with (ii) *RAW 264.7* murine macrophages grown as a two-dimensional (2D) culture. Macrophages, labelled with far-red lysosomal live cell tracker (magenta) consistently integrated with the *AKP* organoids (GFP, green). Macrophages typically migrated to the organoid lumen, suggesting higher self-adhesion (Figure 1a).

**Figure 1:**
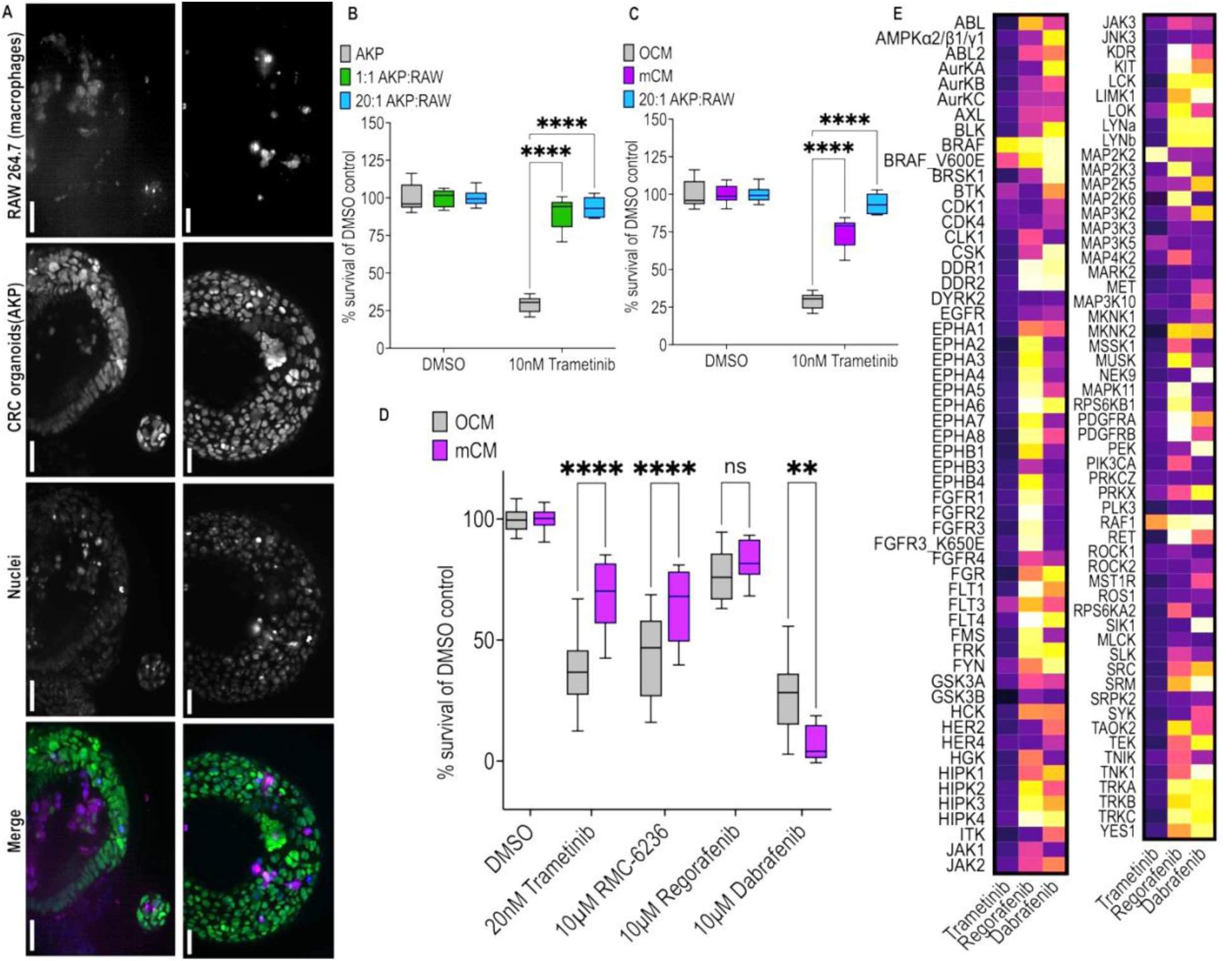
Macrophages drive resistance to trametinib in pre-clinical AKP colorectal cancer organoids. **A)** Representative fluorescence microscopy images showing 48-hour co-culture of *AKP*:*RAW 264.7* mixed at 20:1. *RAW 264.7*, *AKP*, and nuclei are shown in the merged image using magenta, green and blue colour respectively. Both panels show organoids grown in 0.1% DMSO in naïve co-culture media. The scale bar represents 500 µm. **B)** We treated two co-culture ratios with trametinib and measured cell viability. This was compared to trametinib treatment of *AKP* monoculture. **C)** We assessed trametinib response again; in *AKP* we compared viability using OCM or macrophage conditioned media (mCM), and contrasted these to co-culture viability mixed at 20:1 as in panel A. **D)** *AKP* viability in response to treatment with a range of multi-kinase, or targeted, therapies in OCM and mCM. Cell viability was plotted as a box plot. Significance from comparisons is displayed as ****<0.0001, ***<0.001, **<0.01, *<0.05, by two-way ANOVA with Tukey correction. **E)** Heatmap representation of dabrafenib, regorafenib and trametinib’s kinase inhibitory profiles. 100% reflects full loss of kinase activity *in vitro*.

Next, we assessed *AKP* response to the targeted RAS pathway inhibitor trametinib (Mekinist), a potent MEK inhibitor that failed in clinical trials for KRAS mutant CRC [63]. *AKP* organoids were sensitive to treatment with trametinib in naïve organoid culture medium (OCM; Figure 1b). However, co-culture of *AKP* with *RAW 264.7* macrophages, designed to mimic one aspect of a patient’s ITME, resulted in a strong reduction in response to trametinib (Figure 1b). That is, *RAW 264.7* macrophages provide a signal that promoted resistance to at least one potent targeted therapy.

To assess whether this emergent drug resistance was due to a secreted factor, we tested macrophage conditioned media (mCM). mCM alone was sufficient to promote resistance to trametinib: cell viability was increased 260% compared to OCM, though to a lesser extent than the 325% increase in viability observed in *AKP+RAW* co-cultures (Figure 1c). To broaden our work, we assessed the KRAS inhibitor RMC-6236, which is currently in cancer clinical trials (*NCT05379985*). Similar to trametinib, *AKP* cells proved resistant to RMC-6236 when cultured in mCM compared to naïve OCM (Figure 1d). These data suggest that local macrophages provide secreted factors that promote *AKP* resistance to at least two potent RAS-pathway targeted drugs. Data for additional tested drugs is provided below.

Of note, our screens identified two RAS pathway inhibitors that reduced *AKP* survival even in the presence of mCM. Regorafenib, the sole FDA approved multi-kinase inhibitor for KRAS-mutant CRC, showed (limited) ability to kill *AKP* cells at 10 µM; this activity was unaffected by mCM (Figure 1d). Treating *AKP* cells with dabrafenib, used in combination therapies for BRAF mutant colorectal tumours [64–66], led to a stronger reduction in viability; surprisingly, dabrafenib showed still greater potency in mCM to compared to naïve OCM (Figure 1d).

To explore these mCM-associated changes to *AKP* drug sensitivity in response to a range of RAS-pathway inhibitors, we examined previously published *in vitro* inhibition data for trametinib, regorafenib and dabrafenib to assess kinome level effects or patterns. Regorafenib and dabrafenib displayed broad kinase inhibition profiles that contrast with the narrow profile of trametinib (Figure 1e). From this we hypothesised that mCM-induced drug resistance was inhibited by regorafenib and dabrafenib through multi-targeting mechanisms, a testable hypothesis.

### Macrophages secrete PDGF-BB and GDF-15 that together promote trametinib resistance

To identify macrophage-derived resistance-associated secretory factors present in mCM, we used a broad antibody array to capture the presence, or absence, of mouse cytokines/chemokines and ligands. This identified several secreted cytokines, including PDGF-BB and GDF-15, that activate intracellular signalling cascades involving kinases inhibited by regorafenib and dabrafenib (Figure 2a).

**Figure 2:**
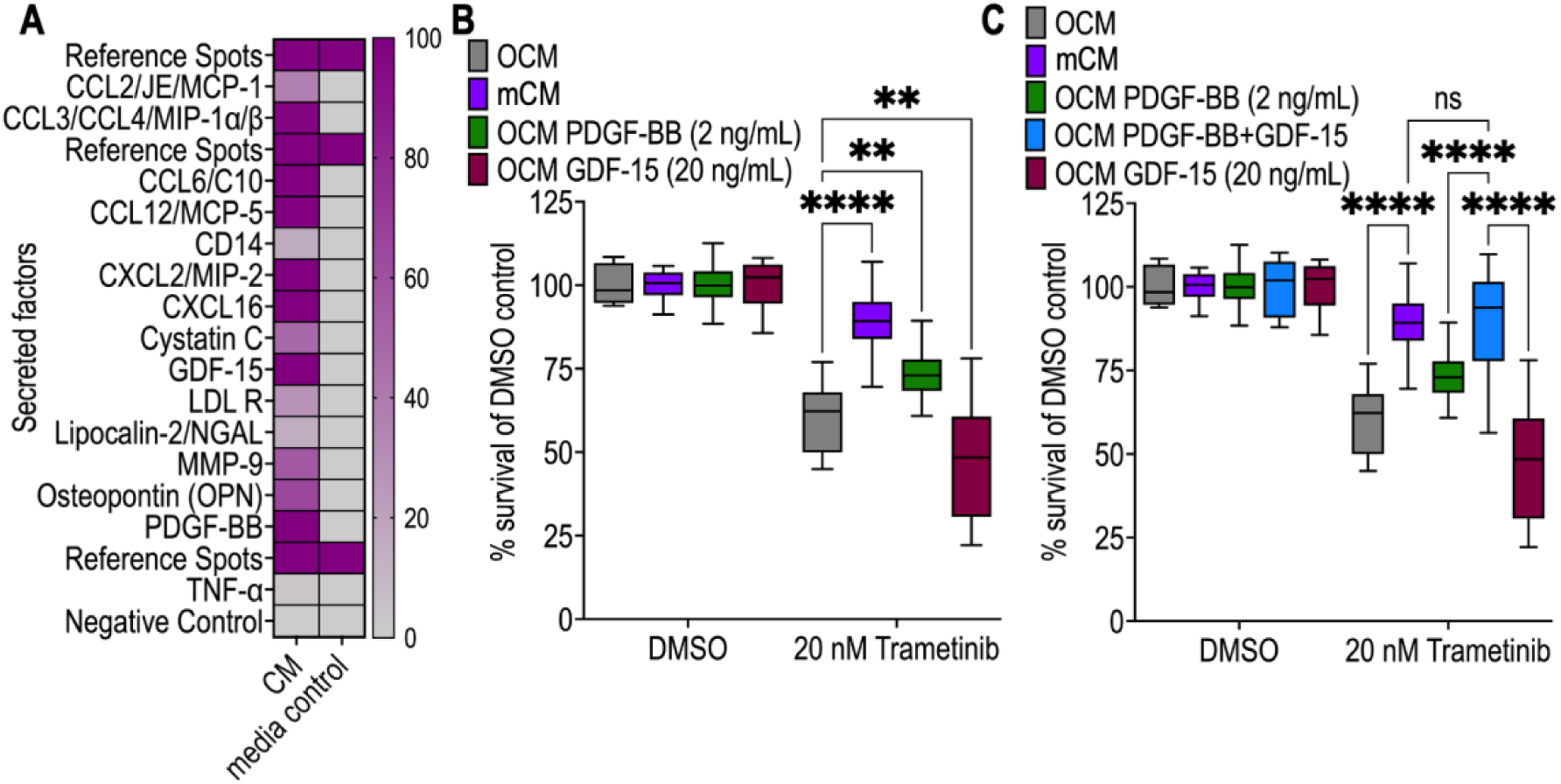
Macrophage-secreted PDGF-BB and GDF-15 alter AKP organoid response to trametinib. **A)** Heatmap showing presence of mouse cytokine and chemokines in conditioned media (mCM) quantified from an antibody array. The percentage intensity is relative to the matched array reference spots. **B)** Selecting hits from the antibody array, we ‘spiked’ recombinant PDGF-BB at 2 ng/mL into OCM and separately doing the same with recombinant GDF-15 at 20 ng/mL. Cell viability was shown in box plot for OCM, mCM and both ‘spiked’ OCM conditions. **C)** Following assessment of recombinant PDGF-BB or GDF-15 impact on trametinib response, we added both recombinant proteins together to OCM at 2 ng/mL and 20 ng/mL respectively. Cell viability was measured for OCM, mCM, OCM+PDGF-BB, OCM+PDGF-BB+GDF-15 and OCM+GDF-15. Significance from comparisons is displayed as ****<0.0001, ***<0.001, **<0.01, *<0.05, by two-way ANOVA with Tukey correction.

To assess the roles of individual ligands, we ‘spiked’ recombinant PDGF-BB or GDF-15 into naïve OCM. Adding recombinant GDF-15 alone decreased *AKP* viability by 20% compared to naïve OCM, indicating increased sensitivity to trametinib. In contrast, recombinant PDGF-BB mildly decreased *AKP* response to trametinib, increasing mean viability by 121%, though 26% less than with mCM (Figure 2b).

Adding both PDGF-BB and GDF-15 into naïve OCM recapitulated levels of mCM-driven trametinib resistance. We show there is no significant difference between mCM viability and ‘spiked’ OCM+PDGF-BB+GDF-15 viability: both increased drug resistance by 147% (Figure 2c). We conclude that mCM-driven resistance to trametinib is multifactorial and likely mediated, at minimum, by PDGF-BB plus GDF-15.

### Macrophages act through RET and PDGFR to promote trametinib resistance

Next, we used targeted inhibitors to examine the roles of the likely GDF-15 and PDGF-BB receptors by comparing their effects on trametinib activity in *AKP* cells cultured in mCM versus OCM.

GDF-15’s receptor complex in the colon has not been characterised; in the brain, RET serves as its primary receptor [54]. Addition of the RET inhibitor GSK3179106 [68] in PDGF-BB+GDF-15 ‘spiked’ media significantly increased *AKP* sensitivity to trametinib, as assessed by cell viability (Figure 3a). This result is consistent with GDF-15 signalling through RET to impact drug response.

**Figure 3.**
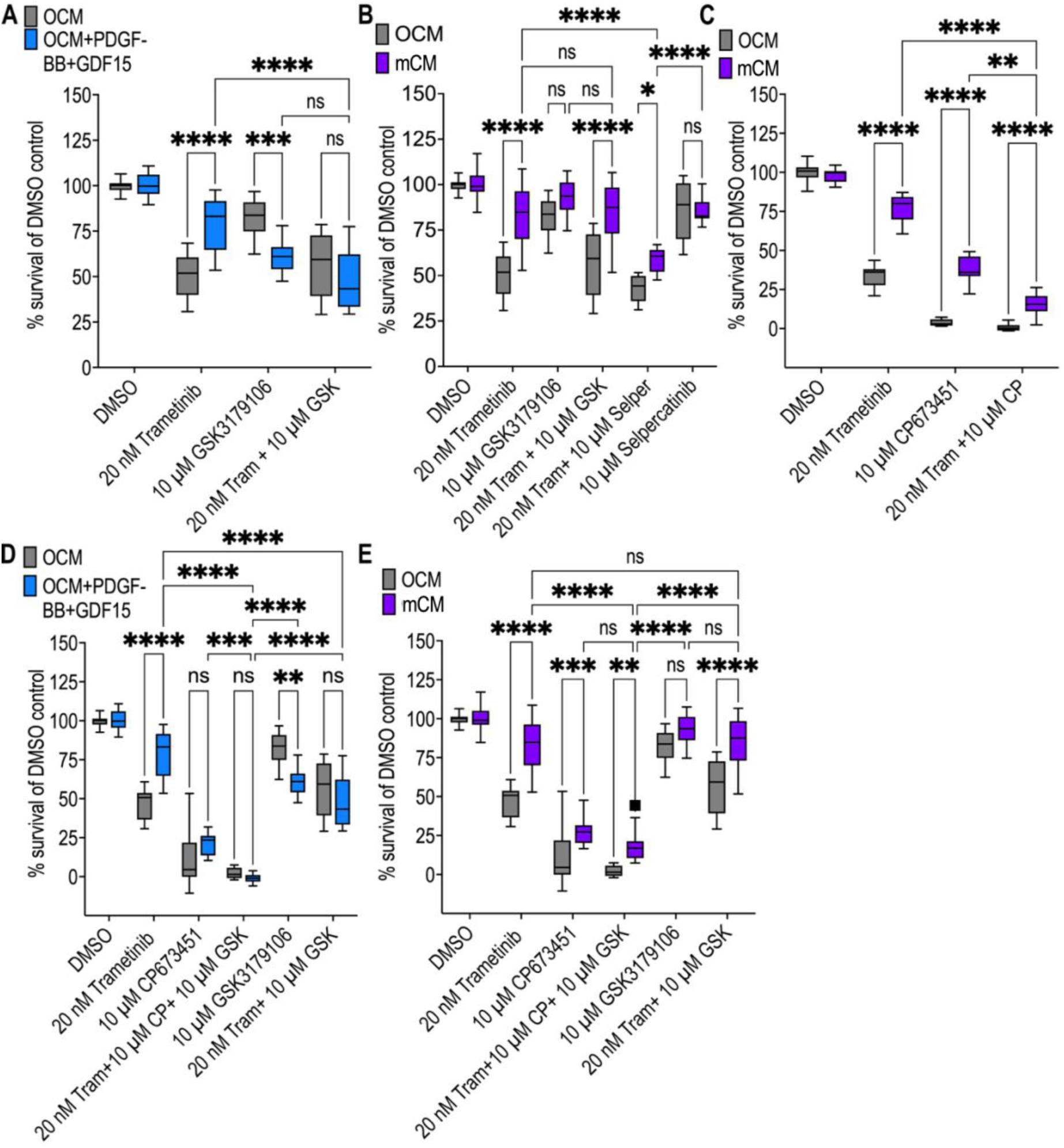
mCM-driven resistance diverges from PDGF-BB+GDF-15-driven resistance. **A)** Viability of AKP organoids cultured in OCM or OCM supplemented with PDGF-BB (2 ng/mL) and GDF-15 (20 ng/mL), following treatment with DMSO, trametinib (20 nM), GSK3179106 (10 µM), or the combination of trametinib and GSK3179106. Survival is shown as a percentage of the matched DMSO control. **B)** Viability of AKP organoids cultured in OCM or mCM following treatment with DMSO, trametinib (20 nM), GSK3179106 (10 µM), selpercatinib (10 µM), or the indicated combinations with trametinib. Survival is shown as a percentage of the matched DMSO control. **C)** Viability of AKP organoids cultured in OCM or mCM following treatment with DMSO, trametinib (20 nM), CP673451 (10 µM), or the combination of trametinib and CP673451. Survival is shown as a percentage of the matched DMSO control. **D)** Viability of AKP organoids cultured in OCM or OCM supplemented with PDGF-BB (2 ng/mL) and GDF-15 (20 ng/mL), following treatment with DMSO, trametinib (20 nM), CP673451 (10 µM), GSK3179106 (10 µM), or the indicated drug combinations, including concurrent MEK, PDGFR, and RET inhibition. Survival is shown as a percentage of the matched DMSO control. **E)** Viability of AKP organoids cultured in OCM or mCM following treatment with DMSO, trametinib (20 nM), CP673451 (10 µM), GSK3179106 (10 µM), or the indicated drug combinations, including concurrent MEK, PDGFR, and RET inhibition. Survival is shown as a percentage of the matched DMSO control. Significance is indicated as ns, *P<0.05, **P<0.01, ***P<0.001, and ****P<0.0001 by two-way ANOVA with Tukey’s multiple-comparison correction.

Following this, we tested AKP response to RET inhibition in mCM using GSK3179106 and multi-kinase RET inhibitor selpercatinib [69]. Both failed to abolish mCM-driven drug resistance, alone or in combination with trametinib (Figure 3b). Additionally, we tested *AKP* response with mCM to the targeted PDGFR inhibitor CP673451 [70]. Inhibiting PDGFR activity with CP673451 failed to reduce mCM-treated response to match OCM-treated response, alone or in combination with trametinib (Figure 3c). We conclude that inhibiting PDGFR or RET alone is not sufficient to eliminate the full mCM-mediated resistance phenotype.

To assess the ability to reduce macrophage-induced drug resistance specifically through PDGFR and RET inhibition, we tested both compounds (CP673451 and GSK3179106 respectively) together with trametinib in a triple combination. We saw *AKP* response to this triple combination, inhibiting PDGFR, RET and MEK, did not vary between ‘spiked’ OCM media and OCM (Figure 3d). However, mCM was still able to drive resistance to this combination, indicating our mCM-driven resistance network includes additional nodes and is mechanistically divergent from ‘spiked’-media driven resistance to trametinib (Figure 3e).

Together, these data suggest that macrophage-derived PDGF-BB and GDF-15 likely contribute to drug resistance through PDGFR- and RET-associated signalling in AKP. Additional macrophage-secreted factors are likely required to explain the full resistance phenotype seen in mCM.

### A role for SRC family members in mediating mCM-driven drug resistance

To identify downstream pathways that mediate mCM-driven resistance, we used PAMGene to compare pathway activities of OCM-*vs.* mCM-treated *AKP* cells. PAMGene technology uses kinase profiling to report on the activities of a broad palette of intracellular pathways.

Under trametinib treatment, mCM increased kinase activity across several signalling networks (Figure 4a). Kinases linked to cell cycle control and JAK/STAT signalling were elevated, with tyrosine kinases showing some of the strongest changes. Notably, PDGFR- and Src family kinase-associated activity were strongly increased, while RET-associated activity was more modestly increased (Figure 4a).

**Figure 4:**
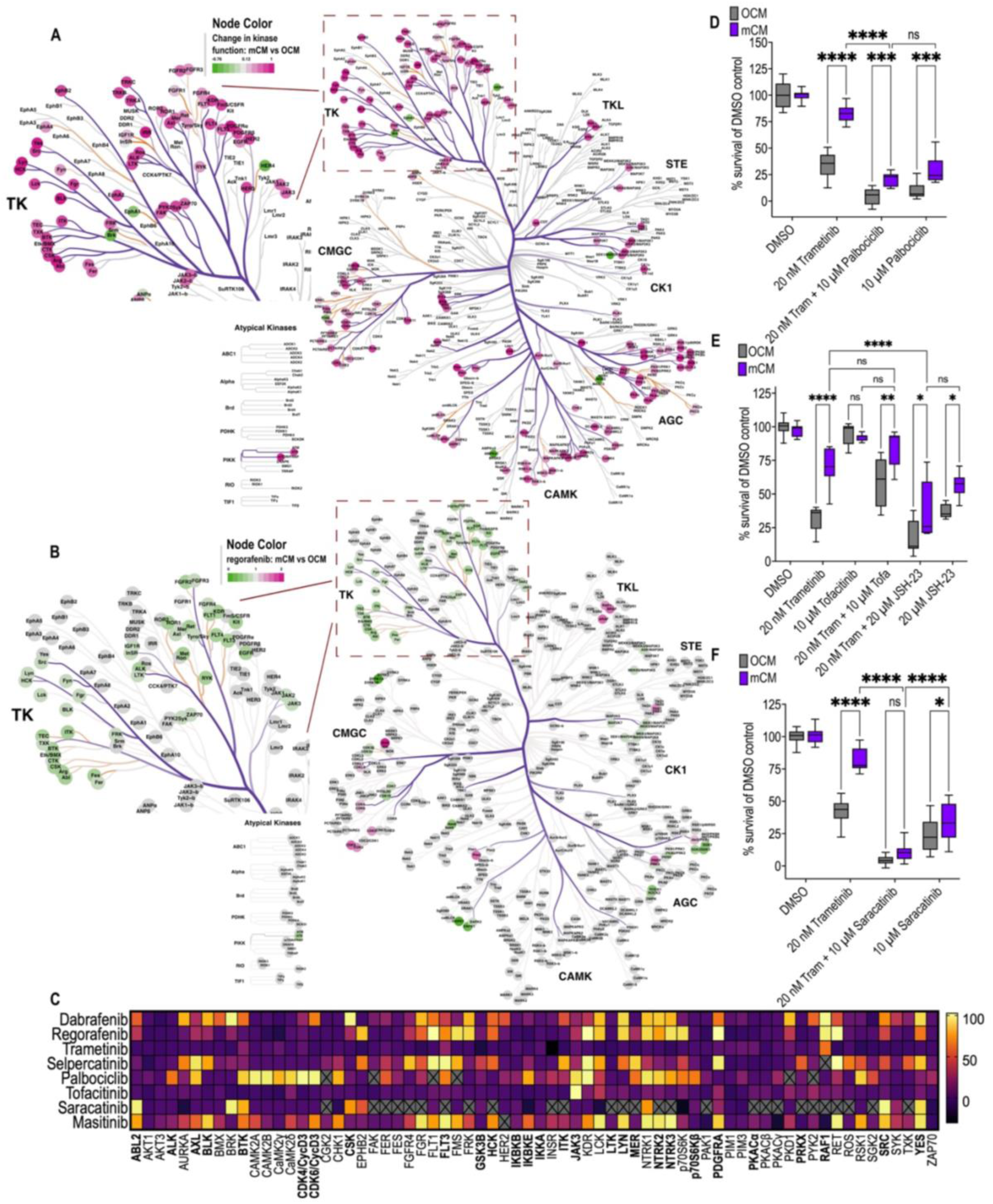
mCM activates Src family kinases to promote trametinib resistance. **A)** Inferred upstream kinase activity in AKP organoids cultured in mCM relative to OCM following treatment with 20 nM trametinib, displayed on the kinome tree using CORAL. Node colour indicates the inferred change in kinase activity in mCM versus OCM, and branch colour indicates the median final score. **B)** Inferred upstream kinase activity in AKP organoids cultured in mCM relative to OCM following treatment with 10 µM regorafenib, displayed on the kinome tree as in panel A. **C)** Heat map showing published in vitro kinase inhibitory profiles of dabrafenib, regorafenib, trametinib, selpercatinib, palbociclib, tofacitinib, saracatinib and masitinib across selected kinases highlighted by the kinome analysis. The colour scale indicates relative inhibition (%); crosses indicate missing data. Most significant hits indicated by **bold**. **D)** Viability of AKP organoids cultured in OCM or mCM following treatment with DMSO, trametinib (20 nM), palbociclib (10 µM), or the combination of trametinib and palbociclib. Survival is shown as a percentage of the matched DMSO control. **E)** Viability of AKP organoids cultured in OCM or mCM following treatment with DMSO, trametinib (20 nM), tofacitinib (10 µM), trametinib plus tofacitinib, JSH-23 (2 µM), or trametinib plus JSH-23. Survival is shown as a percentage of the matched DMSO control. **F)** Viability of AKP organoids cultured in OCM or mCM following treatment with DMSO, trametinib (20 nM), saracatinib (10 µM), or the combination of trametinib and saracatinib. Survival is shown as a percentage of the matched DMSO control. Significance is indicated as ns, *P<0.05, **P<0.01, ***P<0.001, and ****P<0.0001 by two-way ANOVA with Tukey’s multiple-comparison correction.

We previously found that mCM did not functionally alter the response of *AKP* cells to regorafenib. We therefore used PAMGene profiling to compare kinase activity in regorafenib-treated AKP cells cultured in mCM versus OCM. In contrast to trametinib, regorafenib treatment was associated with reduced activity of several tyrosine kinases in mCM, including those associated with PDGFR, RET and SFK signalling (Figure 4b).

Building on these findings, we next asked whether mCM-driven trametinib resistance could be targeted at the network level. We compared published drug inhibitory profiles to kinases activated by mCM from Figure 4a, kinases inhibited by dabrafenib or regorafenib, and kinases not fully covered by selpercatinib (Figure 4c) [71–78]. This allowed us to select compounds to combine with trametinib to target the activated network more broadly.

Based on the increased activity of cell cycle kinases in mCM-treated *AKP* cells under trametinib, we assessed whether these kinases contribute to the drug resistance phenotype. Co-treatment with trametinib and the CDK4/6 inhibitor palbociclib did not reduce mCM-associated drug resistance: viability in mCM was 21% compared to the 4% in OCM (Figure 4d) [79,80]. This suggests that the resistance seen in mCM is not explained simply by increased proliferation.

mCM also increased activity of signalling nodes downstream of receptor tyrosine kinases, including JAK/STAT and NF-κB. To test whether these pathways contribute to trametinib resistance, we combined trametinib with the JAK inhibitor tofacitinib or the NF-κB inhibitor JSH-23 [81,82]. In the presence of either combination, mCM continued to promote trametinib resistance relative to OCM (Figure 4e), indicating that inhibition of either pathway alone was not sufficient to reverse the mCM-driven phenotype.

SFK activity was also increased in mCM-treated AKP cells under trametinib. Because SFKs are key downstream mediators of both PDGFR and RET signalling, we tested saracatinib (AZD0530), a potent inhibitor of SRC and related SFKs, in combination with trametinib [83]. Under this combination, viability fell to 4% in OCM and 10% in mCM (Figure 4f); the difference between the two conditions was no longer significant. This data was consistent with the hypothesis that SFKs are an important downstream component of the mCM-driven resistance network.

### Masitinib combined with MEK or RAS inhibition suppressed macrophage-driven resistance

Our data indicates that macrophages secrete PDGF-BB and GDF-15 and activate intracellular signalling cascades—through canonical receptor activation—that promote resistance to targeted therapies such as trametinib. We therefore used *in vitro* kinase binding data to identify a single multi-targeting drug that targets multiple points along this pathway. Masitinib (AB1010) is a clinically relevant drug that inhibits RET, PDGFRA, cell cycle kinases and multiple SFKs (Figure 5a-b) [84,85].

**Figure 5.**
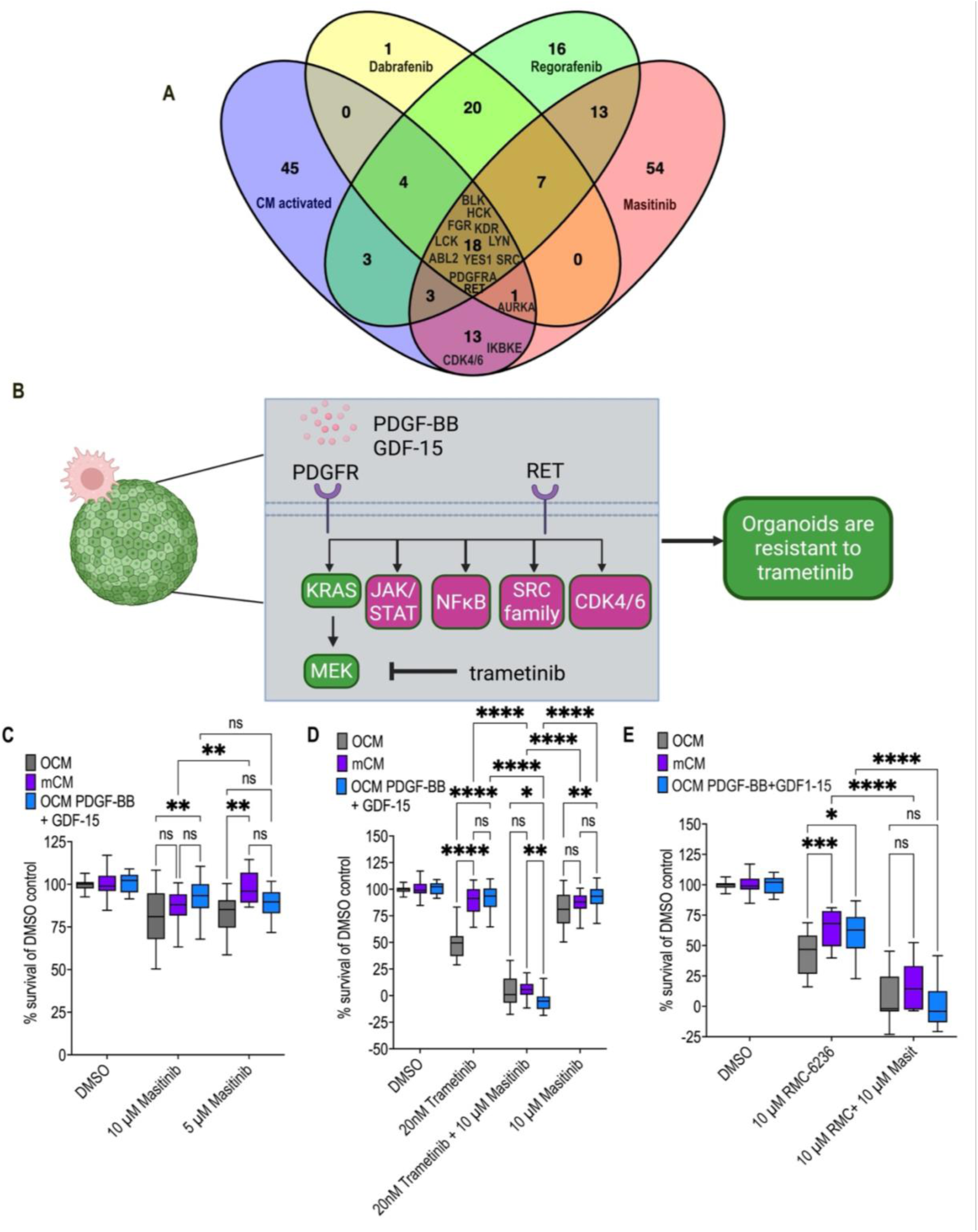
Masitinib suppresses macrophage-driven resistance to trametinib and pan-RAS inhibition. **A)** Venn diagram [88] showing overlap between kinases activated in mCM under trametinib treatment (from Figure 4A) and kinases targeted by dabrafenib, regorafenib, and masitinib (from Figure 4C). Selected shared kinases are indicated. **B)** Schematic model of the proposed macrophage-driven resistance network in which PDGF-BB and GDF-15 signal through PDGFR- and RET-associated pathways to activate downstream nodes including JAK/STAT, NF-κB, SRC family kinases, and CDK4/6, thereby promoting resistance to trametinib. **C)** Viability of AKP organoids cultured in OCM, mCM, or OCM supplemented with PDGF-BB (2 ng/mL) and GDF-15 (20 ng/mL), following treatment with DMSO, 10 µM masitinib, or 5 µM masitinib. Survival is shown as a percentage of the matched DMSO control. **D)** Viability of AKP organoids cultured in OCM, mCM, or OCM supplemented with PDGF-BB (2 ng/mL) and GDF-15 (20 ng/mL), following treatment with DMSO, trametinib (20 nM), masitinib (10 µM), or the combination of trametinib and masitinib. Survival is shown as a percentage of the matched DMSO control. **E)** Viability of AKP organoids cultured in OCM, mCM, or OCM supplemented with PDGF-BB (2 ng/mL) and GDF-15 (20 ng/mL), following treatment with DMSO, RMC-6236 (10 µM), or the combination of RMC-6236 and masitinib (10 µM each). Survival is shown as a percentage of the matched DMSO control. Significance is indicated as ns, *P<0.05, **P<0.01, ***P<0.001, and ****P<0.0001 by two-way ANOVA with Tukey’s multiple-comparison correction.

In *AKP* organoids masitinib had minimal effect on cell viability in OCM, mCM and ‘spiked’ OCM media at 10 µM (Figure 5c). We then used it in combination with trametinib to test whether masitinib could suppress resistance driven by mCM or ‘spiked’ OCM. Masitinib significantly reduced the ability of both mCM and PDGF-BB+GDF-15-‘spiked’ OCM to drive resistance to trametinib (Figure 5d).

Trametinib is a potent MEK inhibitor in the KRAS signalling pathway. The ongoing development of RAS inhibitors pose an opportunity to explore if masitinib is beneficial, in a conserved fashion, to mediate mCM-driven resistance. As shown earlier, mCM decreases sensitivity to RMC-6236 by 20%. This was mirrored in PDGF-BB+GDF-15 ‘spiked’ OCM (Figure 5e) [86]. Importantly, combining masitinib with RMC-6236 abolished resistance driven by PDGF-BB+GDF-15-‘spiked’ OCM and mCM, aligning with our findings using trametinib (Figure 5e). Together, these data support a model in which mCM promotes resistance to both MEK and RAS inhibition through cumulative PDGFR-, RET- and SFK-associated signalling. This resistance phenotype can be manipulated through co-treatment with the multi-kinase inhibitor masitinib.

## Discussion

Targeted therapies for KRAS-mutant colorectal cancer have shown limited success in the clinic; for reasons that remain poorly understood. In this study we focused on drug resistance, mediated by macrophages, which form part of the immune tumour microenvironment in colorectal cancer.

Using *AKP* organoids and macrophage conditioned media, we found that macrophage-derived signals altered the efficacy of multiple targeted drugs including trametinib. Two drugs that were not inhibited by these signals were regorafenib and dabrafenib: overlap in their inhibitory profiles suggested candidate pathways for mediating resistance. Proteomic profiling of conditioned media identified PDGF-BB and GDF-15 as prominent macrophage-secreted factors. Reconstituting naïve organoid media with both recombinant proteins phenocopied resistance to trametinib. Interestingly, GDF-15 alone in naïve media increased *AKP* sensitivity to trametinib. These results support the model that resistance is driven by ligand combinations, not single factors, and that drug resistance is an emergent property of multi-factorial network signalling.

The recombinant PDGF-BB/GDF-15 spike-in experiments allowed us to assess a potential role of RET in GDF-15 activity. Spiked media proved more sensitive to the gut-restricted RET inhibitor GSK3179106 than OCM or mCM, functionally implicating a role of RET activity. Of note, our studies show additional factors outside of PDGF-BB and GDF-15 in mCM promote drug resistance.

Kinome profiling further supported increased PDGFR-associated signalling in mCM-driven resistance to trametinib. We also saw activation of shared downstream effectors, predominantly the SFKs. This supports a model in which (i) PDGF-BB activates PDGFR signalling, (ii) RET signalling mediates GDF-15 activity, and (iii) together they drive resistance in a SFK-dependent manner (Figure 6). This combinatorial model explains why selective inhibition of PDGFR or RET alone did not overcome resistance in mCM.

**Figure 6:**
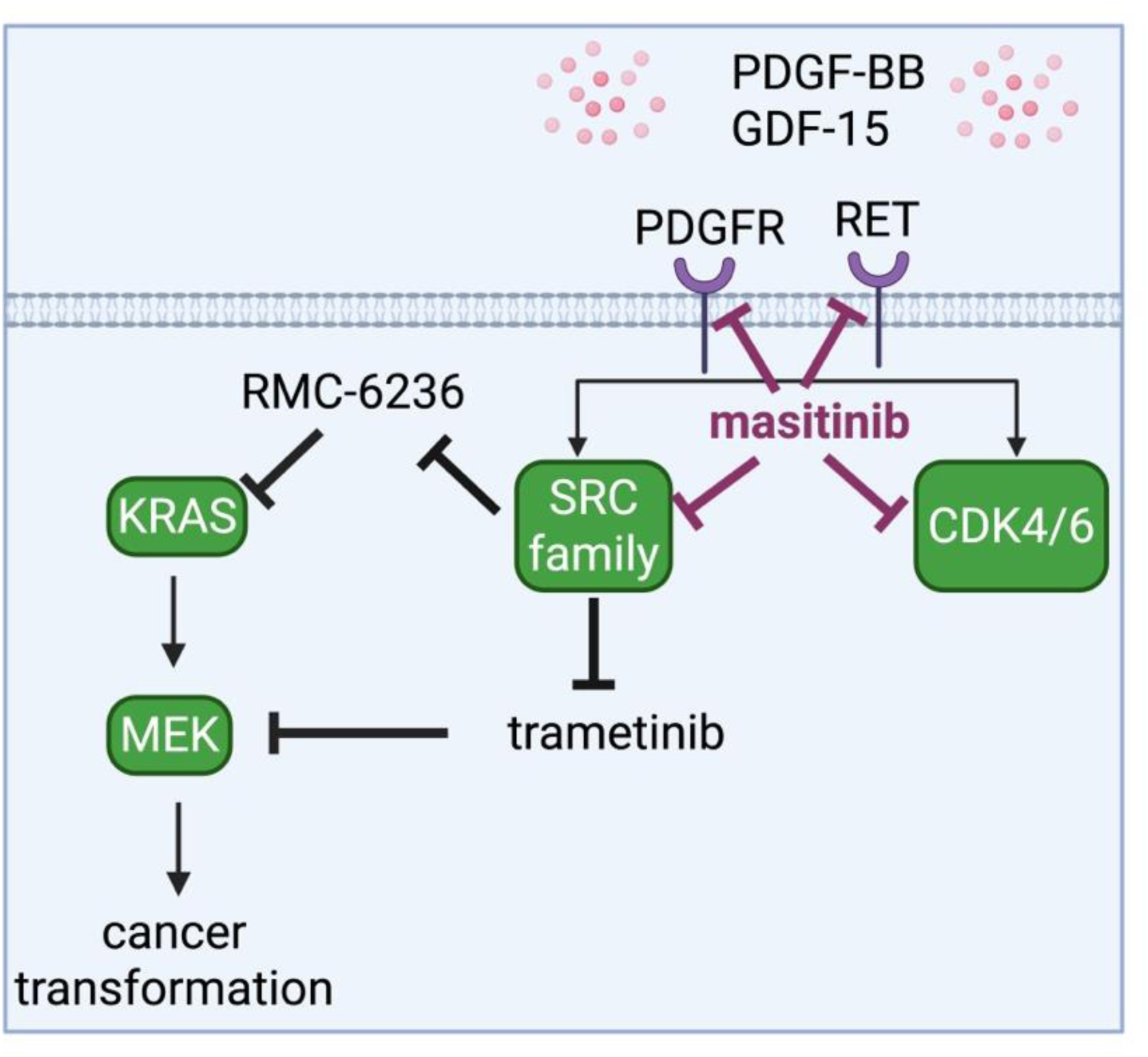
Model of macrophage-driven resistance to MEK and RAS inhibition. PDGF-BB and GDF-15, secreted by the ITME, signal through PDGFR- and RET-associated pathways in KRAS CRC. This results in downstream receptor-associated activation of SRC family kinases (SFKs) and CDK4/6, reducing AKP organoid sensitivity to targeted RAS pathway inhibitors trametinib and RMC-6236. In this model, masitinib concurrently suppresses several key nodes within this network, helping overcome macrophage-conditioned media-driven resistance. BioRender was used to create this model.

These observations support the requirement of a multi-node inhibitory approach rather than a single-target approach. The inhibitory profile of masitinib covers RET, PDGFRs and SFKs, key nodes in our resistance model, among other targets. We leveraged this with RAS pathway inhibitors, trametinib and RMC-6236, to suppress the mCM-driven resistance network.

In summary, we show that macrophage-conditioned media activates resistance networks in our pre-clinical KRAS-mutant colorectal cancer model. We provide evidence that the combination of PDGF-BB and GDF-15 contributes to resistance to targeted therapies. Resistance is associated with increased PDGFR-, RET- and SFK-linked activity, where aligning this network with the inhibitory profiles of multi-kinase inhibitors provides an opportunity to reduce the impact of resistance. More broadly, these findings support a network-informed approach to combination therapy in precision medicine.

## Acknowledgements

We thank the Cagan Laboratory members, Seth Coffelt, Julia Cordero, and Owen Samson for important discussions and supplying *AKP* organoids. This work was generously supported by the Scottish Cancer Foundation and McNab Centre for Cancer Innovation. **R.C.** gratefully acknowledges support from the NIH (R01CA258736), a Royal Society Wolfson Fellowship, Chief Scientific Office (EPD/22/13) (TCS/22/02), CRUK (CTRQQR-2021\100006) Pershing Square Sohn, and Baillie-Gifford. **H.A.B.** gratefully acknowledges support by Tenovus Scotland (S23-01) and a Scotland CSO Fellowship (EPD/23/01).

## Author contributions

**B.S.A.:** conceptualisation, investigation, methodology, validation, formal analysis, visualisation, writing— original draft, writing — review and editing.

**H.A.B.**: supervision, writing — review and editing.

**R.C.**: supervision, funding acquisition, writing — review and editing.

## Competing interests

The authors declare no competing interests.

## Materials and methods

### Murine colorectal cancer organoid culture

Mouse organoid line, AKP, were generated from colony BCVAPK ID:PCB41-1g expressing oncogenic KRAS plus loss of APC and p53 (*Villin–CreERT2: Apc^fl/fl^ Kras^G12D/+^ Trp53^fl/fl^) [87]*, and labelled with green fluorescent protein (GFP) as described below. Derivation of *Vil-Cre^ERT2^*; *Apc^fl/f^*^l^; *Kras^G12D^/+*; *Trp53^fl/fl^* mouse intestinal organoids was performed in accordance with Home Office License: PPL 60/4492, authorised by the University of Glasgow Animal Welfare and Ethical Review Body, and conducted under the UK Animals (Scientific Procedures) Act.

AKP organoids were cultured using Corning® Matrigel® Growth Factor Reduced (GFR) Basement Membrane Matrix, Phenol Red-free, LDEV-free, 10 mL (356231) in Thermo Scientific 6 well Nunc™ Cell-Culture Treated Multidishes (140685). Organoids were cultured in Advanced DMEM + L-Glutamine, Penicillin/streptomycin, HEPES (Advanced DMEM*) supplemented with 50 ng/mL EGF, 100 ng/mL Noggin and 1x N2/B27 (OCM).

Organoids were passaged every 2 to 3 days to maintain healthy structures. Stock organoids in matrigel drops were disrupted in Advanced DMEM and collected into 15ml falcon tubes. Organoid suspension was then centrifuged at 4 °C, 1500 rpm for 8 minutes to separate the Matrigel from the organoid pellet. Media and matrigel were aspirated before further mechanical disruption and resuspension in 6ml of Advanced DMEM. As above, centrifugation and aspiration were repeated, leaving only the organoid pellet. We resuspended cells in 50% Matrigel and seeded them in 20 µL domes.

### Development of labelled organoids

The lentiviral vector *pLV-EKAREN4 (TA)-NLS-Puro* (167830) was obtained from Addgene to label organoids with green fluorescent proteins (GFP). HEK293T cells were obtained from American Type Culture Collection (ATCC). To generate lentiviral particles, HEK293T cells were transfected with 0.5 µg of lentiviral vector, 2 µg of psPAX2 and pCMV-VSV-G (packaging vectors) at 37 °C, 5% CO_2_ for 24 hrs. The transfection supernatant was removed and replaced with organoid culture media. Lentiviral supernatants were collected and cleared using 0.45 µm syringe filter. The supernatant was then used to infect *AKP* organoids in the presence of 10 µg/ml polybrene. GFP organoids were selected using 10 µg/ml of puromycin.

### Murine macrophage culture

*RAW 264.7* cells were obtained from American Type Culture Collection (ATCC). *RAW 264.7* cells were grown in DMEM 10% FBS and 1% Penicillin/streptomycin. Cells were passaged when reaching 70% confluency.

### Drug treatment

Organoids were passaged as above and pellets were then resuspended in a small volume of Advanced DMEM (100-200 µl) for counting. Cell counts were performed using cellometer counting chambers and the cellometer 1000 auto using Trypan blue (Gibco, 11538886) live/dead staining. The live cell count was used to determine the volume of organoid suspension required for experiments. Optical black 96 well plates (PerkinElmer 6005182 Viewplate-96 Black Clear Bottom) were used for drug response assays. *AKP* organoids were seeded at ∼3000 cells/well in a total volume of 6 µL, where the minimum Matrigel content was 50%.

The seeded organoids were treated for 48 hours with the following drugs: trametinib (GSK1120212), regorafenib (BAY 73-4506), dabrafenib (GSK2118436), palbociclib (S4482), JSH-23 (S7351), CP-673451 (S1536), GSK3179106 (S8821), tofacitinib (S2789), selpercatinib (S8781), saracatinib (S1006), masitinib mesylate (E0814), RMC-6236 (E1597) and staurosporine (AM-2282) were obtained from Selleckchem. All compounds were reconstituted according to the manufacturer’s instructions. Staurosporine was used as a positive control at a final concentration of 0.25 µM. Drugs were solubilised in Corning® DMSO (Dimethyl Sulfoxide) (25-950-CQ) for a final dose of 0.1% DMSO.

### Co-culture assays

Organoids and *RAW 264.7* were passaged as above and independently resuspended in media. *AKP* and *RAW 264.7* were mixed in the following ratios: 1:1, 5:1, 10:1, 20:1. The mix of *AKP*:*RAW 264.7* were incubated for 30 minutes at 37 °C. Cells were then resuspended in 80% Matrigel and seeded as 6 µL domes.

### Generation of macrophage conditioned media (mCM)

*RAW 264.7* supernatant was collected before passaging at 70% confluency and centrifuged to remove cells and debris. Supernatant was removed and frozen in aliquots. OCM was mixed prior to experiments in 1:1 ratio with *RAW 264.7* supernatant to create mCM. This mix was then used to perform drug screening.

### Cell Viability Assay

Cell viability was measured using Promega CellTiter-Blue assay kit (G8080) using the manufacturer’s protocol. Plates were loaded into Promega’s Nanoglo machine after 3-4 hours incubation and fluorescence per well was measured at high speed using excitation at 520 nm with emission at 580-640 nm. Measurements were exported into excel files. Three wells per plate were treated with staurosporine to induce 100% cell death. The average fluorescence of the staurosporine positive control wells was subtracted from the plate’s raw data. 100% viability was calculated from the 0.1% DMSO only controls. Each treatment was then calculated as a percentage viability relative to this control. Each media condition was compared and normalised to its corresponding 0.1% DMSO control.

### Fluorescent microscopy

150 nM of Invitrogen lysotracker deep red (L12492) was added directly to *RAW 264.7* culture medium for 1 hour. Media was removed, cells were washed with PBS, and fresh media added again. NucBlue Live ReadyProbes Reagent (R37605) was added as per the manufacturer’s instructions to all cells. Images were obtained using the Opera Phenix and Harmony software. Laser excitation at 405 nm, 488 nm and 640 nm wavelengths was set at 100% power for 100 ms, except for 488 nm at 60 ms. Emission filters 435-480 nm, 500-550 nm and 650-760 nm were used respectively. Fiji was used to analyse and compile the images.

### Secretome profiling

*RAW 264.7* supernatant was collected and screened using Bio-Techne Proteome Profiler Mouse XL Cytokine Array (ARY028). Signal was quantified from chemi-blot images using Fiji and presented as the mean of the two inbuilt replicates.

### Spiked media experiments

Recombinant proteins were purchased from PeproTech: PDGF-BB (315-18) and GDF-15 (120-28C) and diluted as per manufacturer’s instructions. They were diluted in OCM and used for drug screening assays.

### PAMGene Kinase Activity assay

Organoids were treated with 0.1% DMSO, 20 nM trametinib or 10 µM regorafenib for 1 hour before being harvested. All steps were performed on ice. Protocol for 3D organoid lysis was followed as provided by PAMGene published as PROT-7643. An aliquot was taken before storage at −20 °C to perform BCA assays and determine the total protein concentration. Tyrosine and Serine/threonine kinase chips were loaded with lysed samples and data was processed using protocols provided (https://pamgene.com/ps12/) where n=2.

### Statistics and reproducibility

Normalised drug response data was imported into Prism 10 as a grouped analysis. All experiments included a minimum of two variables; type of media (OCM, mCM, ‘spiked’ media) or co-cultures (*AKP*, 1:1 *AKP*:*RAW*, 20:1 *AKP*:*RAW*), and were tested in a minimum of 3 drug conditions (DMSO, single drug, combinations of drugs), requiring a two-way ANOVA. This was performed within Prism to fit a full model with Tukey’s correction for multiple comparisons and the P value was set to 0.05. Normality tests Anderson-Darling and Shapiro-Wilk were performed to test the goodness of fit of the dataset. Row statistics informed the mean cell viability and standard deviation. A minimum of 6 biological replicates were used to generate the graphs, that included a maximum of 3 technical replicates per biological replicate. Biological replicates were defined by separately seeded, treated and assayed plates from passage 4-18 of AKP organoids. Full statistics for each graph are reported in Supplementary table 1.

## Notes

### Competing Interest Statement

The authors have declared no competing interest.

## References

1. Testa U, Pelosi E, Castelli G. Colorectal Cancer: Genetic Abnormalities, Tumor Progression, Tumor Heterogeneity, Clonal Evolution and Tumor-Initiating Cells. Medical Sciences [Internet]. Multidisciplinary Digital Publishing Institute (MDPI); 2018 [cited 2023 Apr 4];6. 10.3390/MEDSCI6020031

2. Zanatto RM, Santos G, Oliveira JC, Pracucho EM, Nunes AJF, Lopes-Filho GJ, et al. Impact of kras mutations in clinical features in colorectal cancer. Arquivos Brasileiros de Cirurgia Digestiva. Colegio Brasileiro de Cirurgia Digestiva; 2020;33:1–5. 10.1590/0102-672020200003E1524

3. Ji J, Wang C, Fakih M. Targeting KRASG12C-Mutated Advanced Colorectal Cancer: Research and Clinical Developments. Onco Targets Ther. Dove Medical Press Ltd; 2022;15:747–56. 10.2147/OTT.S340392

4. Arai H, Battaglin F, Wang J, Lo JH, Soni S, Zhang W, et al. Molecular insight of regorafenib treatment for colorectal cancer. Cancer Treat Rev [Internet]. W.B. Saunders Ltd; 2019 [cited 2025 Jun 26];81:101912. 10.1016/J.CTRV.2019.101912

5. Basso M, Signorelli C, Calegari MA, Lucchetti J, Zurlo IV, Dell’Aquila E, et al. Efficacy of Regorafenib and Trifluridine/Tipiracil According to Extended RAS Evaluation in Advanced Metastatic Colorectal Cancer Patients: A Multicenter Retrospective Analysis. Target Oncol [Internet]. Adis; 2024 [cited 2026 Mar 5];19:371. 10.1007/s11523-024-01050-3

6. Regorafenib for Metastatic Colorectal Cancer: Common Toxicities and Prevention Strategies [cited 2024 May 10]. https://www.cancernetwork.com/view/regorafenib-metastatic-colorectal-cancer.

7. Uitdehaag JCM, De Roos JADM, Van Doornmalen AM, Prinsen MBW, De Man J, Tanizawa Y, et al. Comparison of the Cancer Gene Targeting and Biochemical Selectivities of All Targeted Kinase Inhibitors Approved for Clinical Use. PLoS One [Internet]. Public Library of Science; 2014 [cited 2025 Jun 26];9:e92146. 10.1371/JOURNAL.PONE.0092146

8. Ryan MB, Corcoran RB. Therapeutic strategies to target RAS-mutant cancers. Nat Rev Clin Oncol. Nature Publishing Group; 2018;15:709–20. 10.1038/s41571-018-0105-0

9. Downward J. RAS synthetic lethal screens revisited: still seeking the elusive prize? Clin Cancer Res. American Association for Cancer Research Inc.; 2015;21:1802–9. 10.1158/1078-0432.ccr-14-2180

10. Zhu C, Guan X, Zhang X, Luan X, Song Z, Cheng X, et al. Targeting KRAS mutant cancers: from druggable therapy to drug resistance. Mol Cancer. BioMed Central Ltd; 2022;21. 10.1186/S12943-022-01629-2

11. Zeiser R, Andrlová H, Meiss F. Trametinib (GSK1120212). Recent Results Cancer Res [Internet]. Recent Results Cancer Res; 2018 [cited 2023 Apr 24];211:91–100. 10.1007/978-3-319-91442-8_7

12. Verissimo CS, Overmeer RM, Ponsioen B, Drost J, Mertens S, Verlaan-Klink I, et al. Targeting mutant RAS in patient-derived colorectal cancer organoids by combinatorial drug screening. Elife. eLife Sciences Publications Ltd; 2016;5:e18489. 10.7554/elife.18489

13. Ghosh S, Fan F, Powell RT, Roszik J, Park YS, Stephan C, et al. Vincristine Enhances the Efficacy of MEK Inhibitors in Preclinical Models of KRAS-mutant Colorectal Cancer. Mol Cancer Ther. American Association for Cancer Research Inc.; 2023;22:962–75. 10.1158/1535-7163.MCT-23-0110

14. Bangi E, Ang C, Smibert P, Uzilov A V., Teague AG, Antipin Y, et al. A personalized platform identifies trametinib plus zoledronate for a patient with KRAS-mutant metastatic colorectal cancer. Sci Adv [Internet]. Sci Adv; 2019 [cited 2023 May 5];5. 10.1126/SCIADV.AAV6528

15. Fan F, Ghosh S, Powell R, Roszik J, Park Y, Sobieski M, et al. Combining MEK and SRC inhibitors for treatment of colorectal cancer demonstrate increased efficacy in vitro but not in vivo. PLoS One [Internet]. Public Library of Science; 2023 [cited 2024 Jul 9];18:e0281063. 10.1371/JOURNAL.PONE.0281063

16. Matsubara H, Miyoshi H, Kakizaki F, Morimoto T, Kawada K, Yamamoto T, et al. Efficacious Combination Drug Treatment for Colorectal Cancer That Overcomes Resistance to KRAS G12C Inhibitors. Mol Cancer Ther. American Association for Cancer Research Inc.; 2019;22:529–38. 10.1158/1535-7163.MCT-22-0411

17. Di Veroli GY, Fornari C, Wang D, Mollard S, Bramhall JL, Richards FM, et al. Combenefit: an interactive platform for the analysis and visualization of drug combinations. Bioinformatics. Oxford University Press; 2016;32:2866–8. 10.1093/bioinformatics/btw230

18. Kamal Y, Schmit SL, Frost HR, Amos CI. The tumor microenvironment of colorectal cancer metastases: Opportunities in cancer immunotherapy. Immunotherapy. Future Medicine Ltd.; 2020;12:1083–100. 10.2217/imt-2020-0026

19. Xiang X, Wang J, Lu D, Xu X. Targeting tumor-associated macrophages to synergize tumor immunotherapy. Signal Transduction and Targeted Therapy 2021 6:1 [Internet]. Nature Publishing Group; 2021 [cited 2024 May 10];6:1–12. 10.1038/s41392-021-00484-9

20. Zhang Y, Zhao Y, Li Q, Wang Y. Macrophages, as a Promising Strategy to Targeted Treatment for Colorectal Cancer Metastasis in Tumor Immune Microenvironment. Front Immunol. Frontiers Media S.A.; 2021;12. 10.3389/FIMMU.2021.685978

21. Tan Y, Wang M, Zhang Y, Ge S, Zhong F, Xia G, et al. Tumor-Associated Macrophages: A Potential Target for Cancer Therapy. Front Oncol. Frontiers Media S.A.; 2021;11. 10.3389/FONC.2021.693517

22. Zhu S, Luo Z, Li X, Han X, Shi S, Zhang T. Tumor-associated macrophages: Role in tumorigenesis and immunotherapy implications. J Cancer. Ivyspring International Publisher; 2021;12:54–64. 10.7150/JCA.49692

23. Cao J, Liu C. Mechanistic studies of tumor-associated macrophage immunotherapy. Front Immunol. Frontiers Media SA; 2024;15. 10.3389/FIMMU.2024.1476565

24. Shimizu D, Yuge R, Kitadai Y, Ariyoshi M, Miyamoto R, Hiyama Y, et al. Pexidartinib and Immune Checkpoint Inhibitors Combine to Activate Tumor Immunity in a Murine Colorectal Cancer Model by Depleting M2 Macrophages Differentiated by Cancer-Associated Fibroblasts. Int J Mol Sci. Multidisciplinary Digital Publishing Institute (MDPI); 2024;25. 10.3390/IJMS25137001

25. Xu S, Wang C, Yang L, Wu J, Li M, Xiao P, et al. Targeting immune checkpoints on tumor-associated macrophages in tumor immunotherapy. Front Immunol. Frontiers Media S.A.; 2023;14. 10.3389/FIMMU.2023.1199631

26. Guo S, Chen X, Guo C, Wang W. Tumour-associated macrophages heterogeneity drives resistance to clinical therapy. Expert Rev Mol Med. Cambridge University Press; 2022;24. 10.1017/ERM.2022.8

27. Coletta S, Lonardi S, Sensi F, D’angelo E, Fassan M, Pucciarelli S, et al. Tumor cells and the extracellular matrix dictate the pro-tumoral profile of macrophages in CRC. Cancers (Basel). MDPI; 2021;13. 10.3390/CANCERS13205199

28. Pelka K, Hofree M, Chen JH, Sarkizova S, Pirl JD, Jorgji V, et al. Spatially organized multicellular immune hubs in human colorectal cancer. Cell. Elsevier B.V.; 2021;184:4734–4752.e20. 10.1016/J.CELL.2021.08.003

29. Ammendola M, Curcio S, Ammerata G, Luposella M, Battaglia C, Laface C, et al. Macrophages in Tumor Microenvironment: From Molecular Aspects to Clinical Applications. Eurasian J Med Oncol. Kare Publishing; 2023;7:201–8. 10.14744/EJMO.2023.26480

30. Muller PA, Matheis F, Mucida D. Gut macrophages: key players in intestinal immunity and tissue physiology. Curr Opin Immunol [Internet]. Curr Opin Immunol; 2020 [cited 2024 Jan 19];62:54–61. 10.1016/J.COI.2019.11.011

31. Bain CC, Schridde A. Origin, Differentiation, and Function of Intestinal Macrophages. Front Immunol [Internet]. Front Immunol; 2018 [cited 2024 Jan 19];9. 10.3389/FIMMU.2018.02733

32. Christofides A, Strauss L, Yeo A, Cao C, Charest A, Boussiotis VA. The complex role of tumor-infiltrating macrophages. [cited 2024 Jan 16]; 10.1038/s41590-022-01267-2

33. Yahaya MAF, Lila MAM, Ismail S, Zainol M, Afizan NARNM. Tumour-Associated Macrophages (TAMs) in Colon Cancer and How to Reeducate Them. J Immunol Res [Internet]. J Immunol Res; 2019 [cited 2023 Sep 5];2019. 10.1155/2019/2368249

34. Wang H, Tian T, Zhang J. Tumor-Associated Macrophages (TAMs) in Colorectal Cancer (CRC): From Mechanism to Therapy and Prognosis. Int J Mol Sci [Internet]. Multidisciplinary Digital Publishing Institute (MDPI); 2021 [cited 2022 Nov 7];22. 10.3390/IJMS22168470

35. Shao S, Miao H, Ma W. From Dual Roles to Translational Challenges: Unpacking the Complexities of Tumor-Associated Macrophages in Cancer Progression and Therapy. 2023 [cited 2025 Nov 21]; 10.20944/PREPRINTS202305.1877.V1

36. Bailey C, Negus R, Morris A, Ziprin P, Goldin R, Allavena P, et al. Chemokine expression is associated with the accumulation of tumour associated macrophages (TAMs) and progression in human colorectal cancer. Clin Exp Metastasis [Internet]. Springer; 2007 [cited 2024 Jan 12];24:121–30. 10.1007/S10585-007-9060-3/FIGURES/6

37. Wei C, Yang C, Wang S, Shi D, Zhang C, Lin X, et al. M2 macrophages confer resistance to 5-fluorouracil in colorectal cancer through the activation of CCL22/PI3K/AKT signaling. Onco Targets Ther. Dove Medical Press Ltd; 2019;12:3051–63. 10.2147/OTT.S198126

38. Zhang X, Chen Y, Hao L, Hou A, Chen X, Li Y, et al. Macrophages induce resistance to 5-fluorouracil chemotherapy in colorectal cancer through the release of putrescine. Cancer Lett [Internet]. Cancer Lett; 2016 [cited 2023 Oct 20];381:305–13. 10.1016/J.CANLET.2016.08.004

39. Dadlani E, Dash T, Sahoo D. Investigating tumor-associated macrophages and their polarization in colorectal cancer using Boolean implication networks. 2023 [cited 2025 Nov 21]; 10.1101/2023.08.01.551559

40. Pan Y, Yu Y, Wang X, Zhang T. Tumor-Associated Macrophages in Tumor Immunity. Front Immunol [Internet]. Front Immunol; 2020 [cited 2023 Aug 3];11. 10.3389/FIMMU.2020.583084

41. Hegarty LM, Jones GR, Bain CC. Macrophages in intestinal homeostasis and inflammatory bowel disease. Nature Reviews Gastroenterology & Hepatology 2023 20:8 [Internet]. Nature Publishing Group; 2023 [cited 2024 Jan 16];20:538–53. 10.1038/s41575-023-00769-0

42. Caprara G, Allavena P, Erreni M. Intestinal macrophages at the crossroad between diet, inflammation, and cancer. Int J Mol Sci. MDPI AG; 2020;21:1–31. 10.3390/ijms21144825

43. Liu H, Liang Z, Zhou C, Zeng Z, Wang F, Hu T, et al. Mutant KRAS triggers functional reprogramming of tumor-associated macrophages in colorectal cancer. Signal Transduct Target Ther [Internet]. Signal Transduct Target Ther; 2021 [cited 2023 Sep 5];6. 10.1038/S41392-021-00534-2

44. Li L, Tian Y. The role of metabolic reprogramming of tumor-associated macrophages in shaping the immunosuppressive tumor microenvironment. Biomedicine and Pharmacotherapy. Elsevier Masson s.r.l.; 2023;161. 10.1016/J.BIOPHA.2023.114504

45. Zhu Y, Chen X, Pan Q, Wang Y, Su S, Jiang C, et al. A Comprehensive Proteomics Analysis Reveals a Secretory Path- and Status-Dependent Signature of Exosomes Released from Tumor-Associated Macrophages. J Proteome Res [Internet]. J Proteome Res; 2015 [cited 2024 Jan 12];14:4319–31. 10.1021/ACS.JPROTEOME.5B00770

46. Lungulescu C, Sur D, Răileanu Ștefan, Dumitru Ștefania M, Dumitrescu EA, Lungulescu CV. GDF-15 Signaling Leading to Epithelial-to-Mesenchymal Transition in Colorectal Cancer-a Literature Review. Journal of Medical and Radiation Oncology Journal [Internet]. 2022 [cited 2024 Aug 30];II:1–8. 10.53011/JMRO.2022.01.01

47. Wang D, Townsend LK, DesOrmeaux GJ, Frangos SM, Batchuluun B, Dumont L, et al. GDF15 promotes weight loss by enhancing energy expenditure in muscle. Nature. Nature Research; 2023;619:143–50. 10.1038/S41586-023-06249-4

48. Kim-Muller JY, Song LJ, LaCarubba Paulhus B, Pashos E, Li X, Rinaldi A, et al. GDF15 neutralization restores muscle function and physical performance in a mouse model of cancer cachexia. Cell Rep. Elsevier B.V.; 2023;42. 10.1016/J.CELREP.2022.111947

49. Wang B, Ma N, Zheng X, Li X, Ma X, Hu J, et al. GDF15 Repression Contributes to 5-Fluorouracil Resistance in Human Colon Cancer by Regulating Epithelial-Mesenchymal Transition and Apoptosis. Biomed Res Int. Hindawi Limited; 2020;2020. 10.1155/2020/2826010

50. Zheng H, Yu S, Zhu C, Guo T, Liu F, Xu Y. HIF1α promotes tumor chemoresistance via recruiting GDF15-producing TAMs in colorectal cancer. Exp Cell Res. Academic Press; 2021;398:112394. 10.1016/J.YEXCR.2020.112394

51. Guo Y, Ayers JL, Carter KT, Wang T, Maden SK, Edmond D, et al. Senescence-associated tissue microenvironment promotes colon cancer formation through the secretory factor GDF15. Aging Cell [Internet]. Blackwell Publishing Ltd; 2019 [cited 2025 Dec 2];18:e13013. 10.1111/ACEL.13013

52. Filippini DM, Romaniello D, Carosi F, Fabbri L, Carlini A, Giusti R, et al. The Multifaceted Role of Growth Differentiation Factor 15 (GDF15): A Narrative Review from Cancer Cachexia to Target Therapy. Biomedicines. Multidisciplinary Digital Publishing Institute (MDPI); 2025;13. 10.3390/BIOMEDICINES13081931

53. Ishida J, Konishi M, Saitoh M, Springer J. Growth differentiation factor-15 as a prognostic biomarker in cancer patients. J Cachexia Sarcopenia Muscle. Wiley Blackwell; 2016;7:235–6. 10.1002/JCSM.12123

54. Emmerson PJ, Wang F, Du Y, Liu Q, Pickard RT, Gonciarz MD, et al. The metabolic effects of GDF15 are mediated by the orphan receptor GFRAL. Nat Med. Nature Publishing Group; 2017;23:1215–9. 10.1038/NM.4393

55. Kazlauskas A, Cooper JA. Phosphorylation of the PDGF receptor beta subunit creates a tight binding site for phosphatidylinositol 3 kinase. EMBO J. Springer Science and Business Media LLC; 1990;9:3279–86. 10.1002/j.1460-2075.1990.tb07527.x

56. Heldin CH, Bäckström G, Ostman A, Hammacher A, Rönnstrand L, Rubin K, et al. Binding of different dimeric forms of PDGF to human fibroblasts: evidence for two separate receptor types. EMBO J [Internet]. 1988 [cited 2026 Mar 5];7:1387. 10.1002/j.1460-2075.1988.tb02955.x

57. Morrison DK, Kaplan DR, Rhee SG, Williams LT. Platelet-Derived Growth Factor (PDGF)-Dependent Association of Phospholipase C-γ with the PDGF Receptor Signaling Complex. Mol Cell Biol. Informa UK Limited; 1990;10:2359–66. 10.1128/mcb.10.5.2359-2366.1990

58. Heldin CH, Westermark B. Platelet-derived growth factor: mechanism of action and possible in vivo function. Cell Regul [Internet]. American Society for Cell Biology; 1990 [cited 2026 Mar 5];1:555. 10.1091/mbc.1.8.555

59. Gelderloos JA, Rosenkranz S, Bazenet C, Kazlauskas A. A Role for Src in Signal Relay by the Platelet-derived Growth Factor α Receptor. Journal of Biological Chemistry [Internet]. Elsevier; 1998 [cited 2026 Mar 5];273:5908–15. 10.1074/jbc.273.10.5908

60. Takikita-Suzuki M, Haneda M, Sasahara M, Owada MK, Nakagawa T, Isono M, et al. Activation of Src Kinase in Platelet-Derived Growth Factor-B-Dependent Tubular Regeneration after Acute Ischemic Renal Injury. Am J Pathol [Internet]. American Society for Investigative Pathology Inc.; 2003 [cited 2026 Mar 5];163:277. 10.1016/S0002-9440(10)63651-6

61. Sachsenmaier C, Sadowski HB, Cooper JA. STAT activation by the PDGF receptor requires juxtamembrane phosphorylation sites but not Src tyrosine kinase activation. Oncogene [Internet]. Nature Publishing Group; 1999 [cited 2026 Mar 5];18:3583–92. 10.1038/sj.onc.1202694

62. Shah K, Vincent F. Divergent Roles of c-Src in Controlling Platelet-derived Growth Factor-dependent Signaling in Fibroblasts. American Society for Cell Biology; 2005. 101091/mbc.e05-03-0263

63. Infante JR, Fecher LA, Falchook GS, Nallapareddy S, Gordon MS, Becerra C, et al. Safety, pharmacokinetic, pharmacodynamic, and efficacy data for the oral MEK inhibitor trametinib: A phase 1 dose-escalation trial. Lancet Oncol [Internet]. Elsevier; 2012 [cited 2026 Apr 24];13:773–81. 10.1016/S1470-2045(12)70270-X

64. Corcoran RB, Atreya CE, Falchook GS, Kwak EL, Ryan DP, Bendell JC, et al. Combined BRAF and MEK inhibition with dabrafenib and trametinib in BRAF V600-Mutant colorectal cancer. Journal of Clinical Oncology. American Society of Clinical Oncology; 2015;33:4023–31. 10.1200/JCO.2015.63.2471

65. Salama AKS, Li S, Macrae ER, Park JI, Mitchell EP, Zwiebel JA, et al. Dabrafenib and trametinib in patients with tumors with BRAFV600E mutations: Results of the NCI-MATCH trial subprotocol H. Journal of Clinical Oncology. American Society of Clinical Oncology; 2020;38:3895–904. 10.1200/JCO.20.00762

66. Corcoran RB, Andre T, Atreya CE, Schellens JHM, Yoshino T, Bendell JC, et al. Combined BRAF, EGFR, and MEK inhibition in patients with BRAF(V600E)-mutant colorectal cancer. Cancer Discov. American Association for Cancer Research Inc.; 2018;8:428–43. 10.1158/2159-8290.cd-17-1226

67. Corcoran RB, Atreya CE, Falchook GS, Kwak EL, Ryan DP, Bendell JC, et al. Combined BRAF and MEK inhibition with dabrafenib and trametinib in BRAF V600-Mutant colorectal cancer. Journal of Clinical Oncology. American Society of Clinical Oncology; 2015;33:4023–31. 10.1200/JCO.2015.63.2471

68. Schenck Eidam H, Russell J, Raha K, Demartino M, Qin D, Guan HA, et al. Discovery of a First-in-Class Gut-Restricted RET Kinase Inhibitor as a Clinical Candidate for the Treatment of IBS. ACS Med Chem Lett [Internet]. American Chemical Society; 2018 [cited 2026 Mar 11];9:623. 10.1021/acsmedchemlett.8b00035

69. Bradford D, Larkins E, Mushti SL, Rodriguez L, Skinner AM, Helms WS, et al. FDA Approval Summary: Selpercatinib for the Treatment of Lung and Thyroid Cancers with RET Gene Mutations or Fusions. Clin Cancer Res [Internet]. Clin Cancer Res; 2021 [cited 2026 Mar 11];27:2130–5. 10.1158/1078-0432.CCR-20-3558

70. Roberts WG, Whalen PM, Soderstrom E, Moraski G, Lyssikatos JP, Wang H-F, et al. Antiangiogenic and Antitumor Activity of a Selective PDGFR Tyrosine Kinase Inhibitor, CP-673,451. [cited 2026 Mar 11]; http://aacrjournals.org/cancerres/article-pdf/65/3/957/2538648/957-966.pdf. Accessed 11 Mar 2026

71. Kitagawa D, Yokota K, Gouda M, Narumi Y, Ohmoto H, Nishiwaki E, et al. Activity-based kinase profiling of approved tyrosine kinase inhibitors. Genes Cells [Internet]. Genes Cells; 2013 [cited 2026 Mar 9];18:110–22. 10.1111/gtc.12022

72. Uitdehaag JCM, De Roos JADM, Van Doornmalen AM, Prinsen MBW, De Man J, Tanizawa Y, et al. Comparison of the Cancer Gene Targeting and Biochemical Selectivities of All Targeted Kinase Inhibitors Approved for Clinical Use. PLoS One [Internet]. Public Library of Science; 2014 [cited 2026 Mar 9];9:e92146. 10.1371/journal.pone.0092146

73. Uitdehaag JCM, Kooijman JJ, de Roos JADM, Prinsen MBW, Dylus J, Willemsen-Seegers N, et al. Combined cellular and biochemical profiling to identify predictive drug response biomarkers for kinase inhibitors approved for clinical use between 2013 and 2017. Mol Cancer Ther [Internet]. American Association for Cancer Research Inc.; 2019 [cited 2026 Mar 9];18:470–81. 10.1158/1535-7163.MCT-18-0877

74. Kooijman JJ, van Riel WE, Dylus J, Prinsen MBW, Grobben Y, de Bitter TJJ, et al. Comparative kinase and cancer cell panel profiling of kinase inhibitors approved for clinical use from 2018 to 2020. Front Oncol [Internet]. Frontiers Media S.A.; 2022 [cited 2026 Mar 9];12:953013. 10.3389/fonc.2022.953013

75. Inhibitor | Saracatinib (AZD0530) | International Centre for Kinase Profiling [Internet]. [cited 2026 Mar 9]. https://www.kinase-screen.mrc.ac.uk/inhibitor-and-results/278. Accessed 9 Mar 2026

76. Kim S, Tiedt R, Loo A, Horn T, Delach S, Kovats S, et al. The potent and selective cyclin-dependent kinases 4 and 6 inhibitor ribociclib (LEE011) is a versatile combination partner in preclinical cancer models. Oncotarget [Internet]. Impact Journals LLC; 2018 [cited 2026 Mar 9];9:35226. 10.18632/oncotarget.26215

77. HMS LINCS Project [Internet]. [cited 2026 Mar 9]. https://lincs.hms.harvard.edu/. Accessed 9 Mar 2026

78. Kinase Profiling Inhibitor Database | International Centre for Kinase Profiling [Internet]. [cited 2026 Mar 9]. https://www.kinase-screen.mrc.ac.uk/kinase-inhibitors. Accessed 9 Mar 2026

79. Schmidt M, Sebastian M. Palbociclib-The First of a New Class of Cell Cycle Inhibitors. Recent Results Cancer Res [Internet]. Recent Results Cancer Res; 2018 [cited 2024 May 16];211:153–75. 10.1007/978-3-319-91442-8_11

80. Lee CL, Cremona M, Farrelly A, Workman JA, Kennedy S, Aslam R, et al. Preclinical evaluation of the CDK4/6 inhibitor palbociclib in combination with a PI3K or MEK inhibitor in colorectal cancer. Cancer Biol Ther [Internet]. Cancer Biol Ther; 2023 [cited 2024 May 16];24. 10.1080/15384047.2023.2223388

81. Jiang JK, Ghoreschi K, Deflorian F, Chen Z, Perreira M, Pesu M, et al. Examining the chirality, conformation and selective kinase inhibition of 3-((3R,4R)-4-methyl-3-(methyl(7H-pyrrolo[2,3-d]pyrimidin-4-yl)amino)piperidin-1-yl)-3-oxopropanenitrile (CP-690,550). J Med Chem [Internet]. J Med Chem; 2008 [cited 2026 Mar 11];51:8012–8. 10.1021/jm801142b

82. Shin HM, Kim MH, Kim BH, Jung SH, Kim YS, Park HJ, et al. Inhibitory action of novel aromatic diamine compound on lipopolysaccharide-induced nuclear translocation of NF-κB without affecting IκB degradation. FEBS Lett [Internet]. FEBS Lett; 2004 [cited 2026 Mar 11];571:50–4. 10.1016/j.febslet.2004.06.056

83. Hennequin LF, Allen J, Breed J, Curwen J, Fennell M, Green TP, et al. N-(5-chloro-1,3-benzodioxol-4-yl)-7-[2-(4-methylpiperazin-1-yl)ethoxy]-5-(tetrahydro-2H-pyran-4-yloxy)quinazolin-4-amine, a novel, highly selective, orally available, dual-specific c-Src/Abl kinase inhibitor. J Med Chem [Internet]. J Med Chem; 2006 [cited 2026 Mar 11];49:6465–88. 10.1021/jm060434q

84. Anastassiadis T, Deacon SW, Devarajan K, Ma H, Peterson JR. Comprehensive assay of kinase catalytic activity reveals features of kinase inhibitor selectivity. Nat Biotechnol. 2011;29:1039–45. 10.1038/nbt.2017

85. masitinib | EMD Millipore KinaseProfilerTM screen/Reaction Biology Kinase HotspotSM screen | IUPHAR/BPS Guide to Pharmacology [Internet]. [cited 2026 Mar 9]. https://www.guidetopharmacology.org/GRAC/LigandScreenDisplayForward?ligandId=5656&screenId=3. Accessed 9 Mar 2026

86. Jiang J, Jiang L, Maldonato BJ, Wang Y, Holderfield M, Aronchik I, et al. Translational and Therapeutic Evaluation of RAS-GTP Inhibition by RMC-6236 in RAS-Driven Cancers. Cancer Discov [Internet]. Cancer Discov; 2024 [cited 2026 Mar 11];14:994–1017. 10.1158/2159-8290.CD-24-0027

87. Belmont PJ, Budinska E, Jiang P, Sinnamon MJ, Coffee E, Roper J, et al. Cross-species analysis of genetically engineered mouse models of MAPK-driven colorectal cancer identifies hallmarks of the human disease. DMM Disease Models and Mechanisms [Internet]. Company of Biologists Ltd; 2014 [cited 2026 Mar 9];7:613–23. 10.1242/dmm.013904

88. Venny 2.1.0 [Internet]. [cited 2026 Mar 11].https://bioinfogp.cnb.csic.es/tools/venny/. Accessed 11 Mar 2026

